# SIN3 Regulates Transcriptional and Longevity Responses to Glycolytic Perturbation in *Drosophila melanogaster*

**DOI:** 10.64898/2026.03.09.710676

**Authors:** Anjalie Amarasinghe, Lori A. Pile

## Abstract

Cellular metabolism and gene transcription are closely linked. The conserved transcriptional regulator SIN3 acts as a scaffold for histone deacetylase (HDAC)-containing complexes and is crucial for development, stress resistance, and overall organismal health. SIN3 regulates metabolic gene expression in Drosophila cultured cells, however, an understanding of the extent of its role in coordinating responses to metabolic stress in whole organisms is incomplete. In this study, we explored how SIN3 controls glycolytic gene expression across developmental stages and under genetic and dietary disruption of glycolysis in *Drosophila melanogaster*. Focusing on four key glycolytic enzymes: *phosphofructokinase* (*Pfk*), *enolase* (*Eno*), *pyruvate kinase* (*Pyk*), *and pyruvate dehydrogenase beta* (*Pdhb*), we found that reducing *Sin3A* levels increases their expression in both larvae and adults, indicating that SIN3 plays a consistent role in balancing metabolic gene transcription. Genetic interaction experiments indicate that *Sin3A* interacts with *Pyk* and *Eno*, regulating transcription in a gene-specific manner. Disrupting glycolysis via genetic or dietary means alters glycolytic gene expression, and SIN3 modulates this response. These findings indicate that SIN3 functions as a metabolic sensor, regulating transcription in response to cellular metabolic stress. Additionally, we demonstrate that reducing *Sin3A* levels shortens Drosophila lifespan on both low- and high-sucrose diets, emphasizing the importance of SIN3 in longevity. Overall, these results show that SIN3 is a context-dependent regulator of glycolytic gene expression and lifespan in Drosophila, integrating metabolic signals with chromatin-based transcriptional regulation.

**Summary:** To survive and thrive, organisms must adapt to distinct metabolic inputs. We investigated the response of the conserved transcriptional regulator SIN3 to metabolic stress and its control of glycolytic gene expression in *Drosophila melanogaster*. By measuring glycolytic gene expression, testing genetic interactions, and assessing lifespan under genetic and dietary perturbations, we found that *Sin3A* knockdown elevates glycolytic gene expression in a gene-specific manner and decreases longevity. SIN3 also modulates transcriptional responses to disrupted glycolysis and influences lifespan under sucrose stress. These findings identify SIN3 as a context-dependent transcription regulator that links gene expression with organismal metabolic adaptation.

## Introduction

Gene transcription is a tightly regulated process that enables differential gene expression to align with the dynamic cellular environment. Gene transcription takes place in the chromatin landscape, where DNA is wrapped around histone proteins to form nucleosomes. This nucleosome arrangement can influence the transcription machinery’s access to DNA and, consequently, transcription. Epigenetic modifications on histone tail amino acid residues play a crucial role in regulating transcription (Bannister and Kouzarides 2011). Modifications like acetylation and methylation can either loosen or tighten chromatin structure, thereby affecting how easily the transcriptional machinery accesses the DNA (Kouzarides 2007). These modifications also assist or prevent transcription factors in binding promoter regions (Xin and Rohs 2018; Zhang et al. 2018). Histone-modifying enzyme complexes are responsible for adding or removing these modifications, impacting chromatin accessibility and/or the binding of transcription factors. These reversible, dynamic histone modifications enable cells to precisely modulate gene expression in response to developmental cues, environmental stimuli and cellular stress without changing the DNA sequence.

The relationship between gene transcription and metabolism is a balanced interaction that closely links cellular function with environmental and physiological states. Transcription determines the production of metabolic enzymes and metabolite transporters, shaping the cellular metabolic state. On the other hand, metabolites like acetyl-CoA, S-adenosylmethionine (SAM), and nicotinamide adenine dinucleotide (NAD⁺) act as cofactors or substrates for epigenetic modifications, directly affecting nucleosome arrangement and thereby transcriptional activity (Janke et al. 2015; Yu and Li 2024). This crosstalk between cellular metabolism and transcription regulation ensures that the metabolic state of the cell is closely aligned with gene expression, allowing cells to adjust their transcriptional profile based on nutrient levels, energy status and metabolic stress (Grüning et al. 2010). This reciprocal interaction is vital during development, differentiation, and disease, where metabolic status and epigenetic transcription regulation are equally important (Donati et al. 2018; Carthew 2021). Coordinators of this interaction are not fully understood.

The transcriptional regulator Swi-independent-3 (SIN3) is an evolutionarily conserved transcription co-factor present in many different species, including humans (Nasmyth et al. 1987; Sternberg et al. 1987; Grzenda et al. 2009; Chaubal and Pile 2018). As a scaffold protein, SIN3 recruits multiple protein components to assemble into the SIN3/HDAC co-regulator complex (Grzenda et al. 2009; Adams et al. 2018). SIN3 acts as a soft repressor, reducing expression of many housekeeping genes without completely silencing their expression (Mitra et al. 2021; Soukar et al. 2023). SIN3 is necessary for cell cycle progression, normal development of the organism, and in Drosophila and mammals, the loss of SIN3 results in embryonic lethality (Pennetta and Pauli 1998; Pile et al. 2002; Cowley et al. 2005; Dannenberg et al. 2005; Robert et al. 2023).

SIN3 is essential for the longevity and normal aging of flies and worms, as well as the oxidative stress response (Barnes et al. 2014; Sharma et al. 2018; Konwar et al. 2022; Konwar et al. 2024). SIN3 is essential for regulating cellular metabolism and is required for healthy mitochondrial function and respiration (Pile et al. 2003; Mitra et al. 2022; Giovannetti et al. 2024). SIN3 regulates metabolic enzyme gene expression and a reduction of SIN3 results in changes to metabolite levels in multiple bioenergetic pathways (Liu and Pile 2017; Liu et al. 2020; Konwar et al. 2022). In Drosophila cultured S2 cells, which are derived from embryos (Schneider 1972), SIN3 can detect and respond to changes in cellular metabolic flux by altering gene expression to influence bioenergetics (Soukar et al. 2025). Additionally, loss of SIN3 reduces the β-cell fitness in mice and causes late onset diabetes, with hyperglycemic phenotypes such as reduced circulation insulin and higher fasting blood sugar levels (Yang et al. 2020; Bartolomé et al. 2022). These studies demonstrate the vital role of SIN3 in regulating cellular metabolism. The regulation of gene expression and longevity by SIN3 in response to diet-induced or genetic metabolic stress, however, has not yet been examined.

Here, we investigate the role of SIN3 in the regulation of glycolytic gene expression across different Drosophila growth stages and under various metabolic stress conditions, focusing on four glycolytic genes: *Pfk*, *Eno*, *Pyk* and *Pdhb*. These genes encode enzymes essential for glycolysis and link glycolysis to the tricarboxylic acid (TCA) cycle. *Pfk* and *Pyk* are required for regulatory steps in the glycolysis pathway (Jenkins et al. 2011; Alves-Filho and Pålsson-McDermott 2016). *Eno* is needed for the conversion of 2-phosphoglycerate to phosphoenolpyruvate. *Pdhb* expression links glycolysis to the TCA cycle. SIN3 binding was detected at the promoters of each of these genes (Saha et al. 2016). Our findings demonstrate that *Sin3A* knockdown increases expression of these genes at the larval and adult stages, supporting a model in which SIN3 functions as a regulator to maintain metabolic gene expression throughout development. We found genetic interactions between SIN3 and glycolytic enzymes, and that SIN3 regulates metabolic gene expression when glycolysis is disrupted or metabolic stress is applied. These regulatory effects were also observed to affect organismal longevity under metabolic stress. These findings position SIN3 as a metabolic modulator responsive to genetic and environmental cues, capable of affecting glycolysis and lifespan in Drosophila. Overall, our results establish SIN3 as a context-dependent regulator of glycolytic gene expression and organismal longevity, impacting organismal physiology and stress adaptation.

## Materials and methods

### Fly stocks and husbandry

Fly lines obtained from the Bloomington Drosophila Stock Center are listed in Table 1. The UAS-*Sin3A*^RNAi^ homozygous line and the *Ser*-GAL4> UAS-*Sin3A*^RNAi^ (SIN3 KD II) line were generated in previous studies (Sharma et al. 2008; Swaminathan et al. 2012). The tubulin-GAL4 gene-switch (TGS-GAL4) line (yw; +, tubulin-Gene-Switch) is a gift from Dr. Scott Pletcher (University of Michigan, Ann Arbor, MI). The triple balancer line (w-; Sp/CyO; Sb/TM3-*Ser*) is a gift from Dr. Mark VanBerkum (Wayne State University, Department of Biological Sciences, Detroit, MI). All fly lines were maintained, and fly crosses were performed according to standard laboratory procedures.

**Table 1.** Fly lines obtained from the Bloomington Drosophila Stock Center.

| Stock number | Genotype |
| --- | --- |
| BDSC26300 | y[1] v[1]; P{y[+t7.7] v[+t1.8]=TRiP.JF02070}attP2 |
| BDSC34336 | y[1] sc[*] v[1] sev[21]; P{y[+t7.7] v[+t1.8]=TRiP.HMS01324}attP2 |
| BDSC35218 | y[1] sc[*] v[1]; P{y[+t7.7] v[+t1.8]=TRiP.GL00099}attP2 |
| BDSC9331 | w[1118]; P{w[+mC]=UAS-GFP.dsRNA.R}143 |
| BDSC6791 | w[*]; P{w[+mC]=Ser-GAL4.GF}1 P{Ser-GAL4.GF}2 |
| BDSC1799 | w[*]; P{w[+mC]=GAL4-Hsp70.PB}89-2-1 |
| BDSC20088 | y[1] w[67c23]; P{y[+mDint2] w[+mC]=EPgy2}Pyk[EY10213]/TM3,<br>Sb[1] Ser[1] |

### Heat shock treatment of third instar larvae

Food vials with UAS-*Sin3A*^RNAi^/*HSP70* GAL4 and UAS-*Gfp*^RNAi^/*HSP70* GAL4 third instar larvae (72 h after egg laying) were immersed in a water bath at 37°C for 1 h and, following a 2 h recovery period at 25°C, RNA was extracted using TRIzol^TM^ (Invitrogen) according to the manufacturer’s protocol.

### Gene interaction analysis

Fly crosses and selection were performed as shown in Fig. 2a to obtain double knockdown flies with a genotype of +, +; *Ser*-GAL4>UAS-*Sin3A*^RNAi^, +; UAS-*Pyk*^RNAi^, +. Double knockdown flies and control flies were observed using a dissecting microscope under mild CO_2_ anesthesia. Three replicates were conducted for each cross, and ≥100 flies from each cross were examined. Images of the flies were captured at 30x magnification using a Google Pixel 8a camera mounted on an Olympus SZX16 microscope, and the background was removed in Canva.

### Experimental diets

Fly diets were based on an agar–sugar–yeast medium containing a base of 0.5% (w/v) Bacto™ agar, 8.6% (w/v) propionic acid, 0.06% (v/v) Tegosept (p-hydroxybenzoic acid, methyl ester), Bacto™ yeast extract (w/v) and 5% (w/v) sucrose. To apply dietary stress, all fly food components were kept constant; only the sucrose concentration was changed to either 1% or 20% (w/v). Fly food used in the experiments with TGS-GAL4 gene-switch contained a final concentration of 200 mM RU486 (Sigma-Aldrich) dissolved in 5% ethanol or only 5% ethanol in the control vials.

### Longevity studies

Flies were collected within 24 h after eclosion. Male and female flies were then sorted under mild CO_2_ anesthesia and kept on standard fly food for 48 h at 27°C. Flies were placed on the experimental diets on day three after eclosion. Each vial contained 25 flies. Each replicate included 50 flies for each gender, treatment (either RU486 or ethanol), and for each test and RNAi control. Flies were transferred into new food vials three times a week, and the number of dead flies was recorded each time. The longevity study conducted using different sucrose concentrations was replicated four times, with a total of 200 flies per gender, treatment, test or RNAi control and sucrose concentration. The longevity study of UAS-*Pfk*^RNAi^/TGS-GAL4, UAS-*Eno*^RNAi^/TGS-GAL4, and UAS-*Pyk*^RNAi^/TGS-GAL4 flies and the double knockdown flies (see Supplementary Fig. 2 for the fly crosses) was repeated three times, using a total of 150 flies for each gender, genotype, treatment and for test and RNAi control.

### RNA extraction and quantitative real-time PCR (qRT-PCR)

The same method described in the longevity study section above was used to induce RNAi with RU486 before RNA extraction. Flies placed in different sucrose concentrations were 25 days old when RNA was extracted. Glycolytic gene knockdown flies and the double knockdown flies were 10 days old at the time of RNA extraction due to their shorter lifespan. Heat-shocked larvae were collected after the 2 h recovery period for RNA extraction. Wing imaginal discs were dissected from the wandering third instar larvae. Total RNA was isolated from either adult flies, larvae or larval wing imaginal discs using TRIzol^TM^ (Invitrogen). cDNA was generated from the isolated RNA using the ImProm-II Reverse Transcription System (Promega) using random hexamers (Thermo Fisher Scientific). The cDNA served as the template in the qRT-PCR assay. The analysis was conducted using PowerUp^TM^ SYBR^TM^ Green Master Mix (Thermo Fisher Scientific). Amplification was performed in QuantStudio 3 Real-Time PCR system (Thermo Fisher Scientific). Primers used for analysis are listed in Table 2. Actin was used as the internal control to normalize RNA expression levels.

**Table 2.**
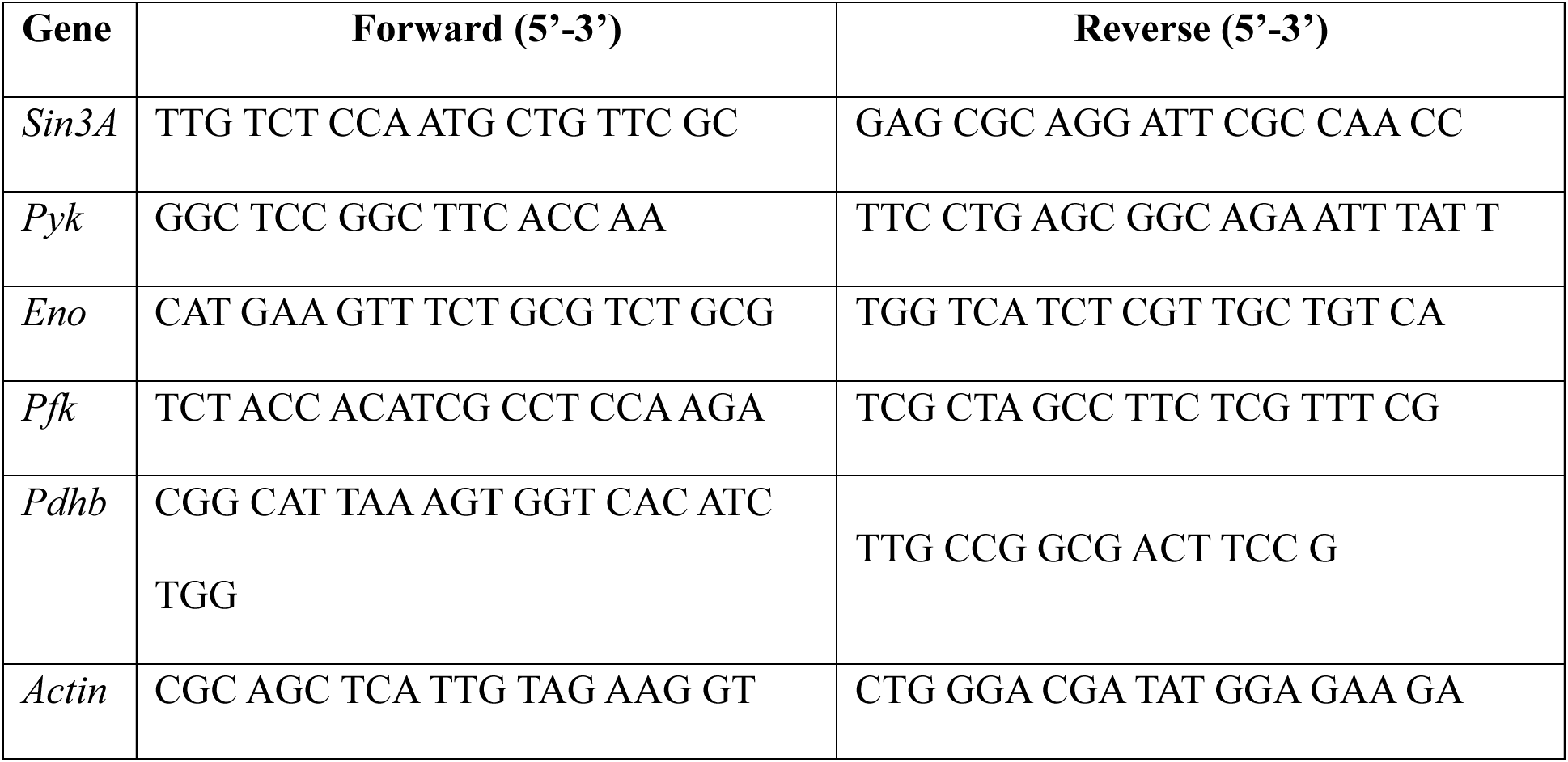
Primers used for the qRT-PCR to amplify the genes analyzed in this study.

### Statistical analysis and gene expression analysis

Prism9 software (GraphPad) was used for all the statistical analyses in this study. A paired Student’s t-test was performed to determine the significance of gene expression level differences. Survival curves were generated using the same software, and Kaplan-Meier survival analysis was performed. The log-rank (Mantel-Cox) test was used for comparing curves.

## Results

### SIN3 regulates expression of *Pfk*, *Eno*, *Pyk* and *Pdhb* in Drosophila third instar larvae and adults

Previous studies indicate that SIN3 regulates the expression of metabolic genes in Drosophila S2 cultured cells (Pile et al. 2003; Saha et al. 2016; Liu et al. 2020). To further investigate this activity, we asked if SIN3 regulates glycolytic gene expression at different stages of Drosophila development. We focused our analysis on the expression levels of the glycolytic genes *Pfk*, *Eno*, *Pyk* and *Pdhb* (Fig. 1a). To assess possible gene regulation by SIN3, we reduced levels of SIN3 through the use of RNA interference (RNAi). To induce RNAi at the larval stage of development, UAS-*Sin3A*^RNAi^ transgenic flies were crossed to the *HSP70* GAL4 driver line. To initiate the RNAi response, third instar larvae produced from that cross were subject to heat shock conditions (37°C for 1h), resulting in ubiquitous GAL4 expression. A control group was maintained at 25°C until RNA extraction was performed. As a control for activation of the RNAi pathway, UAS-*Gfp*^RNAi^ flies were used and crossed to the *HSP70* GAL4 driver line. These UAS-*GfP*^RNAi^ flies have the RNAi machinery activated in their cells, but without a target. *Pfk*, *Eno*, *Pyk* and *Pdhb* RNA levels were significantly upregulated when *Sin3A* was knocked down by approximately 50% (Fig. 1b). We observed that the expression of these genes was slightly reduced in the UAS-*Gfp*^RNAi^ control heat-shocked larvae, consistent with other studies indicating a general down-regulation of gene expression due to the heat treatment (Pessa et al. 2024). To investigate gene expression in adult flies, we used the tubulin gene-switch TGS-GAL4 driver line. In this line, GAL4 is expressed in all tissues upon feeding the flies RU486 (Osterwalder et al. 2001). UAS-*Sin3A*^RNAi^/TGS-GAL4 and UAS-*Gfp*^RNAi^/TGS-GAL4 flies were collected and placed on fly food containing RU486 or ethanol. On day 25 post-eclosion, RNA was extracted for analysis. qRT-PCR analysis showed that adult flies exhibit a similar result to larvae, with the expression of glycolytic genes upregulated in the *Sin3A* knockdown compared to control flies (Fig. 1c). These results demonstrate that, similar to the effect in Drosophila S2 cultured cells, SIN3 regulates glycolytic gene expression in Drosophila larval and adult stages.

**Fig. 1.**
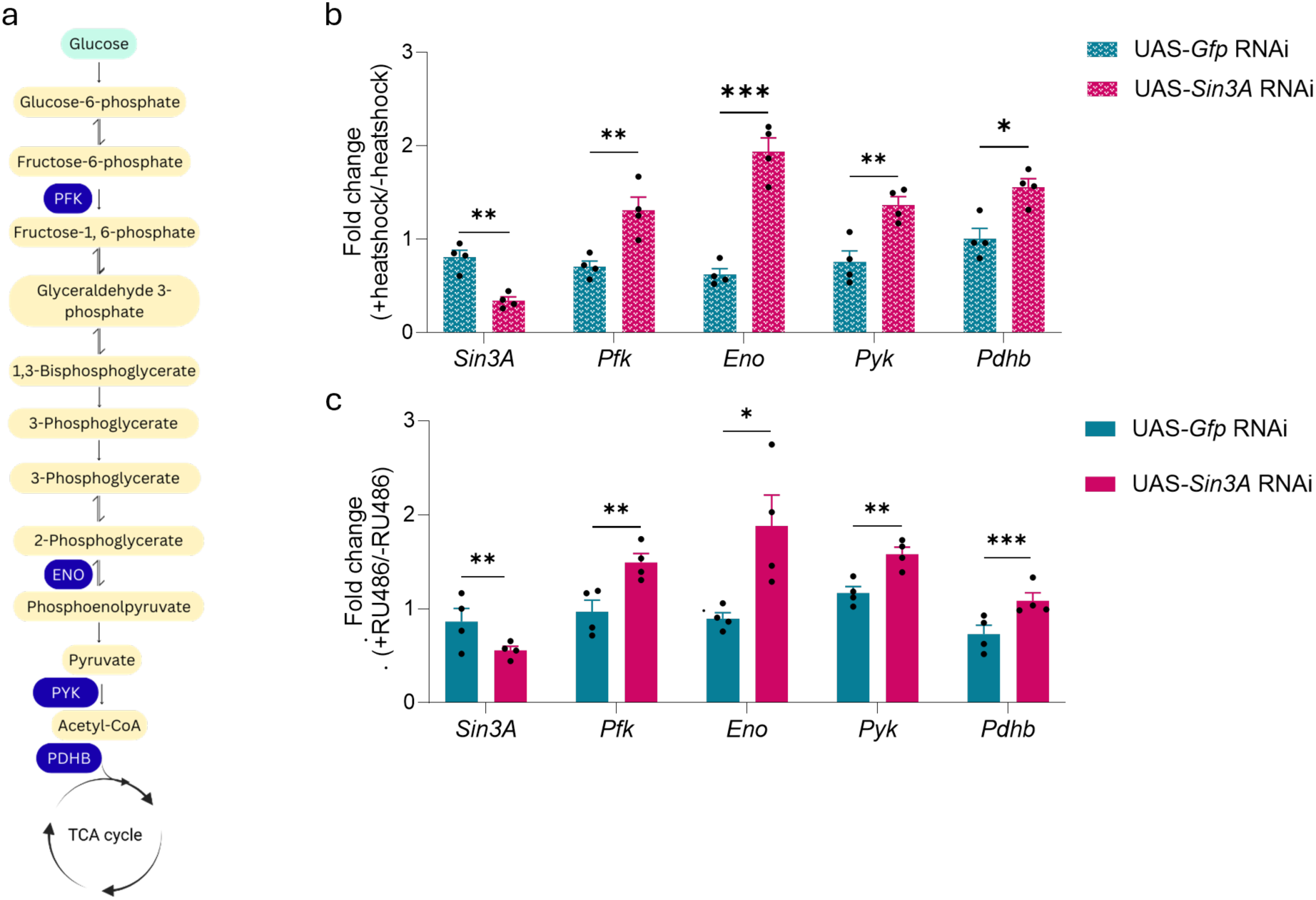
SIN3 regulates gene expression at different developmental stages of Drosophila. (a) Schematic diagram of glycolysis. The enzymes encoded by the genes examined are given in blue. (b) The expression levels of *Eno*, *Pfk*, *Pyk*, and *Pdhb* were compared in larvae for each fly line. Gene expression changes are represented as the mean of the fold changes. Gene expression fold change was calculated as the ratio of UAS-*Gfp*^RNAi^ (heat shock) to UAS-*Gfp*^RNAi^ (room temperature), and of UAS-*Sin3A*^RNAi^ (heat shock) to UAS-*Sin3A*^RNAi^ (room temperature). (c) Gene expression levels of the adult fly experiment that used RU486 to induce RNAi. The fold change calculation of gene expression was made by comparing each fly line treated with RU486 to its respective control reared on standard Drosophila food containing 5% ethanol [e.g., for UAS-*Gfp*^RNAi^, the fold change is given as UAS-*Gfp*^RNAi^(+RU486)/UAS-*Gfp*^RNAi^(ethanol)]. Then, the fold changes for each gene in UAS-*Sin3A*^RNAi^ flies were compared to those in UAS-*Gfp*^RNAi^ flies. \**P* < 0.05, \*\**P* ≤ 0.01, \*\*\**P* ≤ 0.001. Error bars represent standard error of the mean.

### *Sin3A* genetically interacts with *Pyk* and *Eno*

Next, to further analyze the relationship between *Sin3A* and glycolytic genes, we sought to investigate if *Sin3A* has a genetic interaction with *Pyk* and *Eno*. For these studies, we used a fly line previously generated in our laboratory that has *Sin3A* constitutively knocked down by RNAi in wing imaginal discs, *Ser*-GAL4>UAS-*Sin3A*^RNAi^ (Swaminathan et al. 2012). These flies have a curved wing phenotype. To test for a genetic interaction, we generated double knockdown flies and monitored their wings (Fig. 2a). Modification of the curved phenotype with the double knockdown of *Pyk* or *Eno* and *Sin3A* would be indicative of a genetic interaction. *Pyk* and *Sin3A* double-knockdown flies had straight wings with 100% penetrance, indicating a genetic interaction between these genes. We also monitored the wing phenotype of flies with only *Pyk* knocked down in the wing imaginal discs (Fig. 2b). All flies with *Pyk* knockdown had straight wings, indicating that reduction of *Pyk* does not affect wing development. Next, we overexpressed *Pyk* in the wing imaginal discs. Interestingly, these flies had curved wings that phenocopy the *Sin3A* knockdown wing phenotype (Fig, 2b). RNA levels of *Sin3A* and *Pyk* were measured in wing imaginal discs isolated from progeny of the above listed flies (Fig. 2c). RNAi resulted in an approximately 0.6 fold decrease in expression of *Sin3A* and *Pyk* in the respective lines. Consistent with the analysis in whole larvae described above (Fig. 1b), *Pyk* is upregulated approximately 1.8-fold upon reduction of SIN3 by RNAi. *Pyk* knockdown does not affect expression of *Sin3A*.

**Fig. 2.**
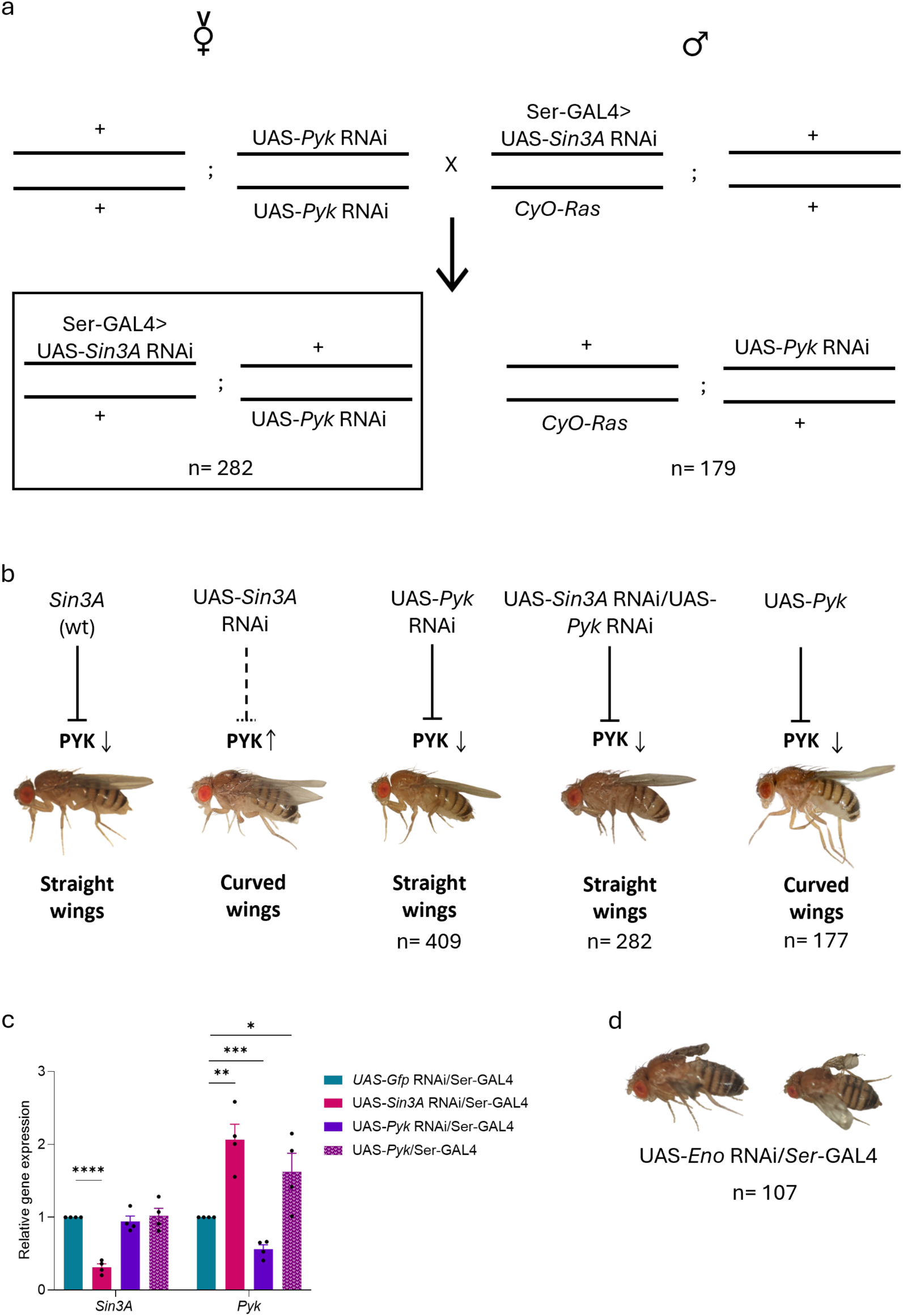
*Pyk* genetically interacts with Drosophila *Sin3A*. (a) Diagram of the fly cross performed using the SIN3 KD II fly line and UAS-*Pyk*^RNAi^. A similar cross was performed between SIN3 KD II and the UAS-*Eno*^RNAi^ line. (b) Images of wing phenotypes observed in each genotype from *Pyk* and *Sin3A* wing imaginal disc knockdown flies and *Pyk* overexpression flies. Female flies are shown. (c) Gene expression analysis showing relative mRNA levels for each genotype shown in b. (d) Female flies with *Eno* knockdown in the wing imaginal discs have blistered wings. \**P* < 0.05, \*\**P* ≤ 0.01, \*\*\**P* ≤ 0.001, \*\*\*\**P* ≤ 0.0001. Error bars represent standard error of the mean.

We also performed a cross between *Ser*-GAL4>UAS-*Sin3A*^RNAi^ and UAS-*Eno*^RNAi^ flies (Supplementary Fig. 1), but did not obtain any viable flies. This result suggested synthetic lethality between *Eno* and *Sin3A.* To determine the impact of reduction of *Eno* alone, we determined the phenotypic effect of *Eno* knockdown in the wing imaginal discs using the *Ser*-GAL4 driver line. All the female progeny from the cross of *Ser*-GAL4 with UAS-*Eno*^RNAi^ had blistered wings (Fig. 2d). We did not obtain any male flies with *Eno* knockdown in the wing imaginal discs. The finding that single knockdown of *Eno* and dual knockdown of *Eno* and *Sin3A* led to lethality was unexpected given the use of the *Ser*-GAL4 driver. We note, however, that *serrate* expression has been detected in tissues other than wing imaginal discs (Chintapalli et al. 2007; Graveley 2010). Together, these results indicate *Eno* is essential for male viability and normal wing development of female flies and that dual reduction in levels of *Sin3A* and *Eno* is synthetically lethal for Drosophila females.

### SIN3 is required to maintain *Pfk*, *Eno*, *Pyk* and *Pdhb* expression when glycolysis is disrupted

With the evidence showing a strong interaction between SIN3 and glycolytic genes, we next asked whether SIN3 is involved in regulation of glycolytic gene expression when the glycolytic pathway is disrupted at different steps. To address this question, we generated fly lines that have the UAS-*Sin3A*^RNAi^ transgene and one of the UAS-*Pfk*^RNAi^, UAS-*Eno*^RNAi^, or UAS-*Pyk*^RNAi^ transgenes. These flies were crossed to the TGS-GAL4 driver line. These fly lines enabled us to induce RNAi machinery in adult flies by adding RU486 to the food, resulting in double knockdown flies (Supplementary Fig. 1). When *Pfk* was knocked down, expression of *Eno*, *Pyk* and *Pdhb* was significantly reduced relative to control flies (Fig. 3a). Simultaneous knockdown of *Sin3A* resulted in upregulation of the expression of these three genes, similar to the effect due to reduction of *Sin3A* alone. This finding suggests that SIN3 is needed to repress the expression of *Eno*, *Pyk* and *Pdhb* in response to glycolysis disruption at the PFK-regulated step. When *Eno* was knocked down, we observed a significant reduction in *Pyk* compared to controls (Fig. 3b)*. Pdhb* levels trended lower. Upon dual knockdown of *Eno* and *Sin3A*, there was an increase in both *Pyk* and *Pdhb* expression relative to the control flies. This finding indicates that SIN3 is necessary for the reduced expression of *Pyk* and *Pdhb* following *Eno* disruption. *Pyk* knockdown caused a significant increase in *Pfk* expression and a decrease in *Pdhb*. This result is consistent with a previous study where the researchers observed an increase in *Pfk* expression following *Pyk* knockdown larvae in an RNA sequencing (RNA-seq) experiment (Heidarian et al. 2024). With *Sin3A* knockdown, *Pfk* levels remained high, and *Pdhb* levels increased significantly compared to either single knockdown of *Pyk* and control lines (Fig. 3c), suggesting that SIN3 is required for *Pdhb* regulation when *Pyk* is reduced. Overall, these results show that SIN3 is involved in regulating a gene-specific response to glycolysis disruption.

**Fig. 3.**
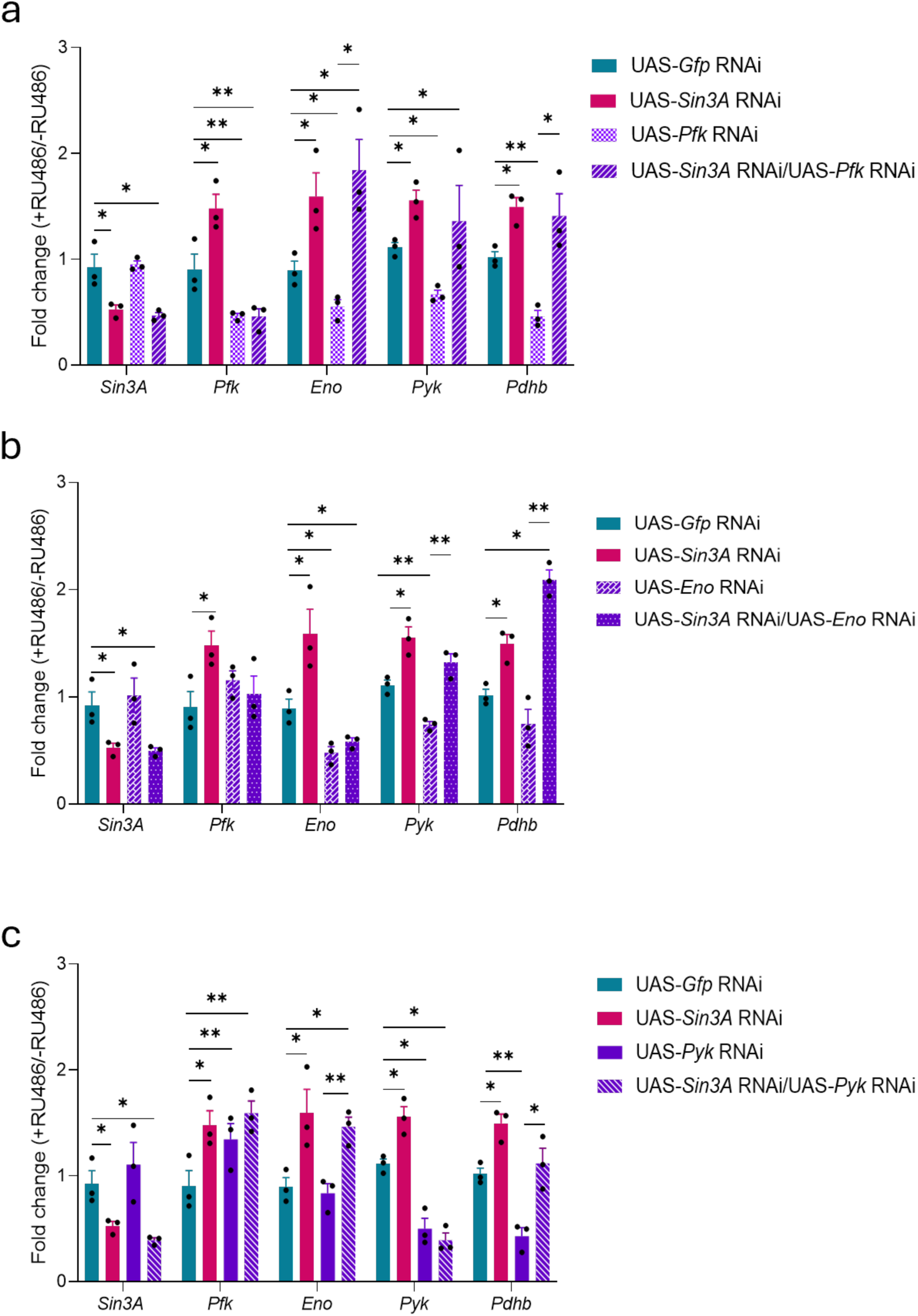
SIN3 is required for the regulation of glycolytic genes when glycolysis is disrupted. (a–c) Gene expression analysis for each knockdown fly line. All flies were placed in vials containing RU468 or 5% ethanol-based food on day 3 post-eclosion. From the glycolytic gene knockdown lines and the double knockdown lines, RNA was extracted for gene expression analysis on day 10. \**P* < 0.05, \*\**P* ≤ 0.01. Error bars represent standard error of the mean.

### Reduced expression of *Sin3A* partially rescues the reduction in Drosophila longevity due to *Pfk* or *Pyk* knockdown

A previous study indicates that induced glycolysis through overexpression of the glycolytic enzymes *triosephosphate isomerase* and *phosphoglucose isomerase* increases Drosophila longevity (Ma et al. 2018). Furthermore, reduced levels of genes encoding glycolytic enzymes *Pyk*, *Pfk* and *Pdhb* results in shortened lifespan (Dung et al. 2018; Wong et al. 2019; Hunt and Demontis 2022; Heidarian et al. 2024) or neuronal and glial cell damage in the case of *Pyk* and *Pdhb* (Volkenhoff et al. 2015; Dung et al. 2018; Waller et al. 2025). Here, we examined how disrupting glycolysis by knocking down *Pfk*, *Eno* or *Pyk* in combination with reduction of SIN3 impacts Drosophila lifespan. We conducted a longevity study with Drosophila subjected to individual knockdown of *Sin3A*, *Pfk*, *Eno* or *Pyk*, as well as double knockdowns with *Sin3A*. *Pfk* knockdown flies had a short lifespan (mean=26.56 days) compared to the mean lifespan of 71.2 days for control flies (Fig. 4a, b). Consistent with our previous study, *Sin3A* knockdown flies had a shortened lifespan, with a mean lifespan of 43.98 days (Barnes et al. 2014). When *Sin3A* was knocked down simultaneously with *Pfk*, the lifespan reduction due to RNAi of *Pfk* was partially rescued, approaching that observed in *Sin3A*^RNAi^ flies. Knockdown of *Eno* drastically reduced fly lifespan (mean=22.32 days), and double knockdown with *Sin3A* resulted in a further decline (Fig. 34, d), consistent with the synthetic lethality seen in the genetic interaction experiment described above (Fig. 2d). *Pyk* knockdown produced results similar to *Pfk* knockdown; double knockdown with *Sin3A* partially rescued reduced fly longevity due to PYK reduction (Fig 4e, f). Female and male flies did not show any statistically significant differences in longevity. Survival curves shown here are for female flies and the data for male flies are given in Supplementary Fig. 2. These results demonstrate that the loss of *Pfk* or *Pyk* reduces Drosophila longevity, and the simultaneous reduction of SIN3 levels can partially rescue this phenotype. Dual reduction of *Eno* and *Sin3A* results in a synergistic decrease in longevity, consistent with the observed synthetic lethality using the *Ser*-GAL4 driver line to induce RNAi.

**Fig. 4.**
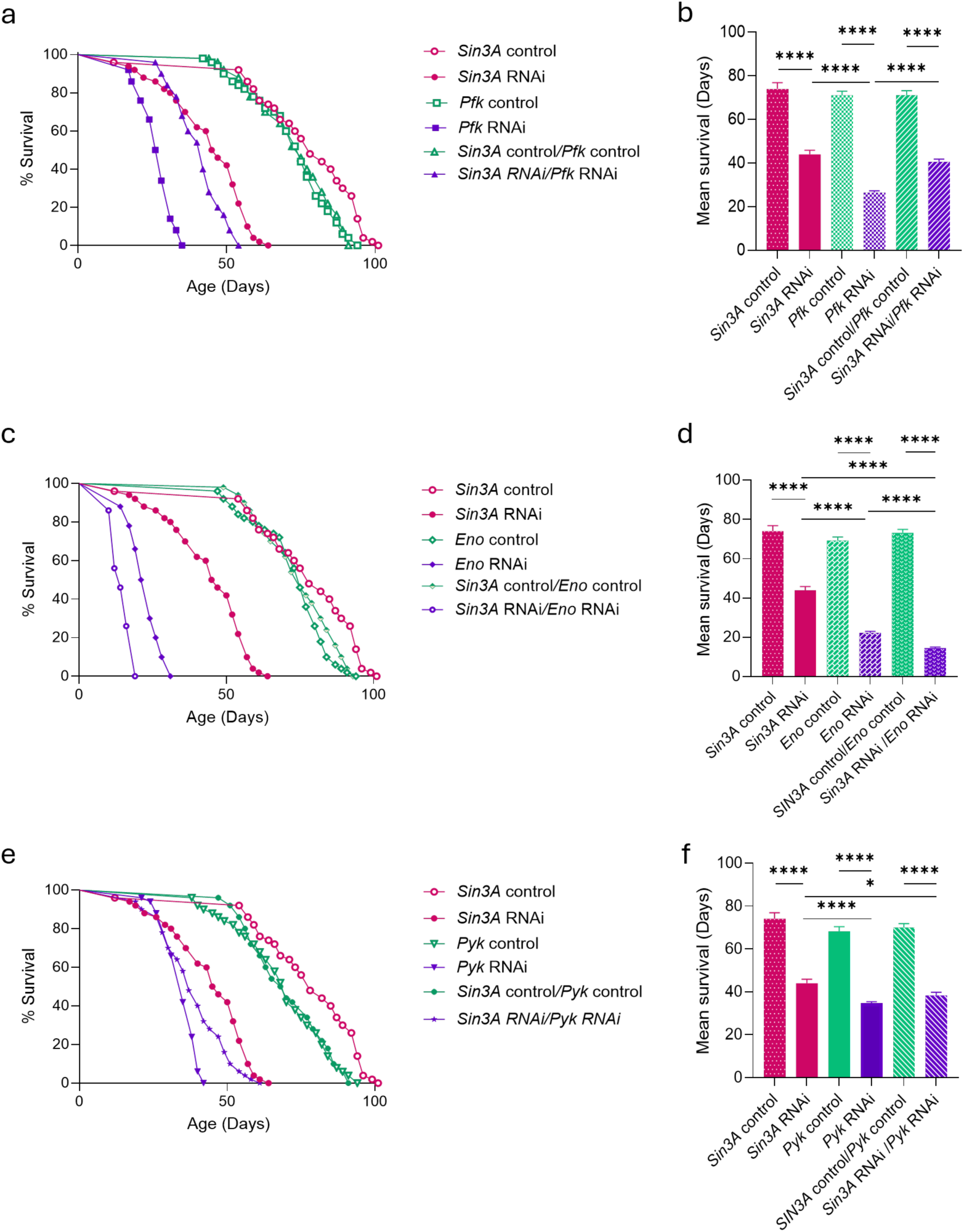
Knocking down *Sin3A* partially rescues the *Pyk* and *Pfk* knockdown phenotypes but not the *Eno* knockdown phenotype. (a,c,e) Survival curves of adult Drosophila for the following conditions: *Sin3A* knockdown alone, *Pyk*, *Eno*, or *Pfk* knockdown alone, and simultaneous knockdown of *Sin3A* with *Pyk*, *Eno*, or *Pfk*. (b,d,f) Mean survival of each genotype. \**P* < 0.05, \*\**P* ≤ 0.01, \*\*\**P* ≤ 0.001, \*\*\*\**P* ≤ 0. 0001. Error bars represent standard error of the mean.

### *Sin3A* and dietary sucrose stress have an additive effect on adult Drosophila longevity

Dietary sucrose concentration can influence Drosophila longevity. Both high and low extreme sucrose concentrations in fly food result in a significant loss of longevity (Magwere et al. 2004; Chandegra et al. 2017; Strilbytska et al. 2020). SIN3 also affects adult Drosophila longevity, with a reduction in levels leading to decreased lifespan (Fig. 4) (Barnes et al. 2014). Therefore, we aimed to examine the impact of SIN3 on Drosophila lifespan when flies are subjected to dietary stress by varying sucrose levels in the food source. For these experiments, we conducted a longevity study on UAS-*Sin3A*^RNAi^/TGS-GAL4 and UAS-*Gfp*^RNAi^/TGS-GAL4 flies maintained on different sucrose concentrations in their food. We used 1% and 20% sucrose concentrations as the two extreme conditions, and standard fly food with 5% sucrose (van Dam et al. 2020; Strilbytska et al. 2020). We observed a significant decrease in longevity in control flies fed 1% sucrose (Fig. 5a, b) compared to those fed 5% sucrose, consistent with previous results (Jans et al. 2024). When *Sin3A* was knocked down and flies were fed 1% sucrose, they exhibited the lowest longevity (mean= 26.5 days). With 20% sucrose, we observed a reduction in lifespan in the control flies (mean= 61.98 days) (Fig. 5c, d). This finding aligns with observations from previous studies (Rovenko et al. 2015; Chandegra et al. 2017; van Dam et al. 2020). Similar to the finding with 1% sucrose, flies with reduced level of SIN3 maintained on 20% glucose had lower longevity compared to controls. We did not observe any statistical significance between male and female Drosophila longevity. Data for female flies are shown in Fig. 5 and the survival curves for male flies are given in Supplementary Figs. 3 and 4. Together, these results demonstrate that SIN3 expression is necessary to maintain Drosophila longevity under two extreme dietary sucrose conditions; the negative impact on longevity caused by 1% and 20% sucrose foods is enhanced by the loss of SIN3.

**Fig. 5.**
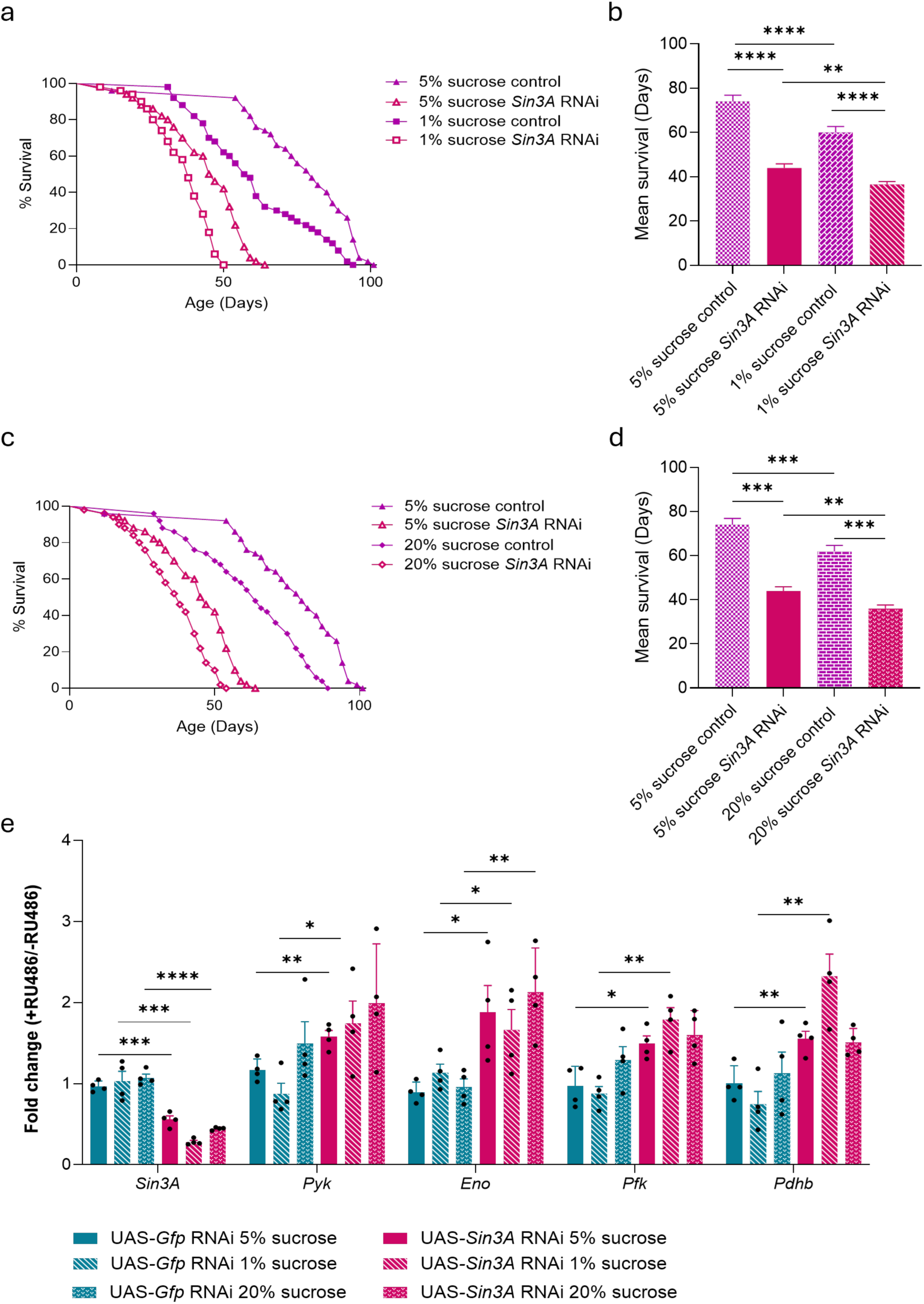
SIN3 and the sucrose stress have an additive effect on Drosophila longevity. (a, c) Survival curves of adult Drosophila reared on food containing 1%, 5%, or 20% (w/v) sucrose. (b, d) Bar graphs showing mean survival. (e) Gene expression analysis of four glycolytic genes under different sucrose concentrations used in the survival analysis. SIN3 is required to maintain *Pyk*, *Eno*, *Pfk* and *Pdhb* expression under different sucrose stress conditions. \**P* < 0.05, \*\**P* ≤ 0.01, \*\*\**P* ≤ 0.001, \*\*\*\**P* ≤ 0. 0001. Error bars represent standard error of the mean.

### *Sin3A* expression is required to repress *Pfk*, *Eno*, *Pyk* and *Pdhb* expression under dietary sucrose stress conditions

Next, we investigated possible alterations to glycolytic gene expression by SIN3 in response to dietary sucrose stress. We examined gene expression in flies experiencing dietary stress. When comparing the gene expression of the selected glycolytic genes in UAS-*GfP*^RNAi^ control flies grown in the 1% and 20% extreme sucrose concentrations to the same genotype grown in standard fly food, we did not observe any significant changes in RNA levels (Fig. 5fe. Next, we compared the UAS-*Gfp*^RNAi^ controls to UAS-*Sin3A*^RNAi^ reared in the different sucrose concentrations independently. In standard fly food conditions, when compared to UAS-*Gfp*^RNAi^ controls, all four glycolytic genes tested were overexpressed in the *Sin3A* knockdown flies (Fig. 5e). A similar result was observed in the UAS-*Sin3A*^RNAi^ flies cultured on 1% sucrose, glycolytic gene expression increased relative to controls (Fig. 5e). In 20% sucrose, only *Eno* showed a significant increase in gene expression in the UAS-*Sin3A*^RNAi^ flies compared to the UAS-*Gfp*^RNAi^ control, though *Pfk*, *Pyk* and *Pdhb*, exhibited a consistent trend toward up-regulation of gene expression following reduction of *Sin3A*. Overall, these findings suggest that SIN3 acts as a key repressor of glycolytic gene expression in standard conditions and under low sucrose stress, with its regulatory influence diminished under high sucrose stress.

## Discussion

In this study, we investigated the impact of the SIN3 co-regulator on glycolytic gene expression across various developmental stages in Drosophila, and under different metabolic stress conditions. The increase in *Pfk*, *Eno*, *Pyk* and *Pdhb* expression following *Sin3A* knockdown in both larvae and adults highlights the role of SIN3 as a transcriptional repressor of key glycolytic genes, consistent with the previous findings in S2 cells (Pile et al. 2003; Saha et al. 2016; Liu et al. 2020).

*Sin3A* genetically interacts with a large and diverse group of genes in Drosophila (Swaminathan et al. 2012). Here, we found that *Sin3A* interacts with *Pyk* and *Eno*. The double knockdown of *Sin3A* and *Pyk* in wing imaginal discs rescued the curved wing phenotype caused by *Sin3A* knockdown alone. This interaction indicates that *Pyk* acts downstream of *Sin3A*. Overexpression of *Pyk* in the wing imaginal discs phenocopies the *Sin3A* knockdown phenotype, further confirming this interaction. The synthetic lethality observed with dual *Sin3A* and *Eno* knockdown shows that *Sin3A* supports compensatory or protective pathways that buffer against the loss of *Eno*. *Eno* depletion alone disrupted wing development in the female flies and caused lethality in males. Although the exact reason for the male-specific lethality observed following *Eno* knockdown in wing imaginal discs is unclear, we suspect that the lethality in males may be due to the knockdown of *Eno* in other tissues, resulting from *Ser* expression in areas other than wing imaginal discs.

The downstream effects of reducing levels of glycolytic enzymes highlight key regulatory steps within the pathway. *Pfk* knockdown led to decreased expression of *Eno*, *Pyk* and *Pdhb*, which encode proteins downstream of Pfk in glycolysis. The simultaneous knockdown of *Sin3A* led to their overexpression. This reversal of the reduced expression following *Pfk* knockdown demonstrates a feedback mechanism mediated by SIN3. Our findings support the model that flux through PFK-mediated steps precisely controls downstream glycolytic enzymes and helps coordinate glycolysis and transition to the TCA cycle (Grüning et al. 2010; Tanner et al. 2018). Conversely, *Eno* knockdown did not significantly alter the expression of glycolytic genes upstream of *Eno*. Still, *Eno* knockdown reduced the downstream *Pyk* expression and we saw the same trend of downregulation in *Pdhb* expression. This response required the presence of SIN3, as SIN3 reduction resulted in increased *Pyk* and *Pdhb* expression in the *Eno*, *Sin3A* double knockdown relative to both control files and to those with *Eno* knockdown alone. There may be a buffering mechanism in place to compensate for the overall effect on glycolysis caused by the loss of *Eno*. However, this suggestion requires further analysis (Poyatos 2020). These gene-specific changes suggest that SIN3 functions as a regulator to balance glycolytic pathway activity and maintain energy homeostasis after disruption of glycolysis following altering levels of enzymes in the pathway.

We observed that *Pyk* knockdown led to increased expression of *Pfk*, which is well upstream of PYK in the glycolytic pathway. A previous RNA-seq experiment in *Pyk* knockdown Drosophila larvae revealed that *Pfk* was overexpressed (Heidarian et al. 2024). Decreasing levels of pyruvate increase PFK activity; however, this occurs through allosteric activators of the enzyme such as low adenosine triphosphate (ATP) levels. (Nunes et al. 2016; Locasale 2018). The increase in *Pfk* transcription following *Pyk* knockdown is more of a gradual adaptation. One possibility is that cells detect reduced glycolytic flux and low ATP production, leading to compensatory upregulation of transcription of the upstream glycolytic enzymes, with PFK regulating a key control point. Another possibility is that the increased PFK enzymatic activity due to low ATP levels leads to rapid conversion of fructose-6-phosphate to fructose-1,6-bisphosphate (Wang et al. 2024), thereby mimicking increased glycolytic flux at this step and signaling the cell to increase *Pfk* transcription. *Pyk* knockdown did not alter *Eno* expression, indicating that disrupting glycolysis specifically affects the upstream rate-limiting step. The knockdown of *Pyk,* however, led to reduced *Pdhb* levels, and this reduction was not observed when *Sin3A* was knocked down simultaneously. *Pdhb* transcribes a subunit in the pyruvate dehydrogenase complex (PDC), which catalyzes the conversion of pyruvate into acetyl-CoA that enters the TCA cycle for producing mitochondrial ATP through oxidative phosphorylation. In Drosophila larvae, knockdown of *Pyk* did not reduce the level of pyruvate (Heidarian et al. 2024). Drosophila larvae rely on glycolysis for energy metabolism (Tennessen et al. 2014). Therefore, other compensatory mechanisms are expected to keep the larval pyruvate levels in check (Heidarian et al. 2024). In the adult flies we examined, the significant reduction of *Pdhb* when *Pyk* is knocked down may lead to reduced levels of pyruvate available to enter the TCA cycle. Metabolomic analysis will be needed to ascertain the outcome.

Previous studies have demonstrated that reducing the expression of specific glycolytic genes reduces organismal overall fitness and longevity, consistent with our results. In mice, mutation of the muscle-specific *Pfk (Pfkm)* gene causes hemolysis, leading to cardiac hypertrophy with age and a reduced lifespan (García et al. 2009). Although Drosophila lacks a *Pfk* muscle-specific gene, we cannot exclude the possibility that comparable effects occur in Drosophila. *Pfk* knockdown causes lethality in Drosophila larvae, indicative of growth and development defects (Wong et al. 2019). These adverse effects of *Pfk* knockdown that reduce the fitness, growth and development of the organism may be the reason for the decreased lifespan measured following *Pfk* knockdown in adult flies observed in this study. Reduced *Pfk* expression in the brain neurons is associated with aging in Drosophila (Oka et al. 2021). Severe locomotion defects were noted in adult Drosophila following the knockdown of *Eno* in panglial cells, and lethality resulted following knockdown of *Pyk* in panglial cells and motor neurons (Volkenhoff et al. 2015; Waller et al. 2025). This disruption of glycolysis in neurons may decrease overall longevity, which may explain the shortened lifespan we observed in *Eno* or *Pyk* knockdown flies. Another study conducted in Drosophila larvae found that knocking down *Eno* in adipose cells significantly reduced glycolysis and decreased adipose cell size (Rodríguez-Vázquez et al. 2024). Furthermore, the authors found downregulation of the mTOR pathway and REPTOR transcriptional activity in Drosophila larvae, leading to muscle disorganization. Neuronal-specific knockdown of *Pdhb* in Drosophila leads to neuronal abnormalities and reduced lifespan (Dung et al. 2018). In our study, *Pdhb* levels were significantly reduced in *Pfk* and *Pyk* knockdown flies, contributing to reduced longevity. We observed that *Sin3A* knockdown partially rescues the reduced lifespan associated with *Pfk* and *Pyk* depletion. This effect may result from transcriptional compensation that lifts the *Sin3A* suppression of essential glycolytic and mitochondrial genes to support cellular energy needs. Conversely, dual reduction of *Eno* and *Sin3A* exacerbated mortality, in line with the wing imaginal disc *Eno* knockdown data. Drosophila ENO, which is similar to mammalian ENOα and ENOγ isoforms (Bishop and Corces 1990), may carry out moonlighting activities that are unrelated to the glycolytic enzyme function. For example, ENOα acts as a transcription regulator (Díaz-Ramos et al. 2012). These activities, which are still not fully investigated, can complicate the overall effects we observe with the *Eno* knockdown.

The reduction of SIN3 is associated with reduced longevity and low stress tolerance in Drosophila (Barnes et al. 2014) and *C. elegans* (Pandey et al. 2018; Sharma et al. 2018; Konwar et al. 2022; Giovannetti et al. 2024; Konwar et al. 2024). In this current study, we saw that the *Sin3A* knockdown enhanced the effect of dietary stress on longevity. Drosophila longevity decreased under both extreme conditions: low (1% w/v) and high (20% w/v) sucrose diets, as reported in previous studies (Lushchak et al. 2014; Strilbytska et al. 2020). The low sucrose diet is associated with oxidative stress (Rovenko et al. 2015). Reduction of *Sin3A* decreases cellular resistance to reactive oxygen species in both flies and worms (Barnes et al. 2014; Sharma et al. 2018). Thus, the summative effects on oxidative stress may lead to the additive effect on longevity we observed in Drosophila cultured on 1% sucrose and having reduced SIN3 levels. High sucrose consumption is associated with obesity and insulin resistance in Drosophila (Rovenko et al. 2015; van Dam et al. 2020). In mice, SIN3 is found as an essential transcriptional regulator that maintains β-cell fitness and thereby prevents diabetes (Yang et al. 2020; Bartolomé et al. 2022). With the high-sucrose diet, we observed an additive effect on reduced Drosophila lifespan with SIN3 reduction, that may be due to the exacerbation of the adverse effects due to the dietary stressor. The transcriptional changes caused by genetic perturbation of key glycolytic enzymes were more pronounced than those observed under dietary sucrose stress, highlighting the importance of gene regulatory mechanisms compared to nutritional factors in maintaining the glycolytic transcriptome. The transcription of the glycolytic genes we investigated did not change significantly in response to altered sucrose levels alone, indicating that SIN3 is able to modulate glycolytic gene expression regardless of sugar availability. These data suggest that transcriptional regulation of these glycolytic genes by SIN3 is independent of two extreme sucrose levels in the diet. A genome-wide analysis is necessary to fully understand the extent of the role of SIN3 in regulating Drosophila longevity under these dietary stress conditions.

Taken together, these results suggest that SIN3 functions as a metabolic mediator, dynamically regulating glycolytic gene expression in a cellular metabolic condition- and gene-dependent manner. The repressive activity of SIN3 maintains transcriptional output under normal conditions but can be lifted to accommodate metabolic disturbances. While dietary sucrose affects lifespan, our data highlight that direct disruption of glycolytic enzymes and their coordinated regulation by SIN3 significantly impacts organismal physiology and gene expression.

## Conclusions

Our findings establish SIN3 as a key regulator of glycolytic gene expression and metabolic adaptation across growth stages in Drosophila. By combining genetic and dietary changes, we demonstrate that SIN3 controls transcription throughout the glycolytic pathway and influences the response to metabolic stress. Gene-specific interactions reveal that SIN3 fine-tunes metabolic output in a context-dependent manner. The reduced longevity of *Sin3A* knockdown flies demonstrates that SIN3 links transcriptional control with whole-organism physiology. Although dietary sucrose stress influences longevity, transcriptional responses to glycolytic disruption are primarily driven by genetic perturbation rather than by nutrient availability. These findings position SIN3 as a key regulator of gene expression that aligns gene expression with cellular energy needs, highlighting its broader role in maintaining metabolic balance and overall organism health.

## Data availability

Drosophila strains are available upon request. The authors affirm that all the data necessary for confirming the conclusions of the article are present within the article, figures, tables, and supplementary files.

Supplemental material available at *GENETICS* online.

## Acknowledgements

We thank current and former members from Dr. Pile’s laboratory and Dr. Sokol Todi for critical evaluation of the manuscript.

---

**Supplementary Fig.1.**
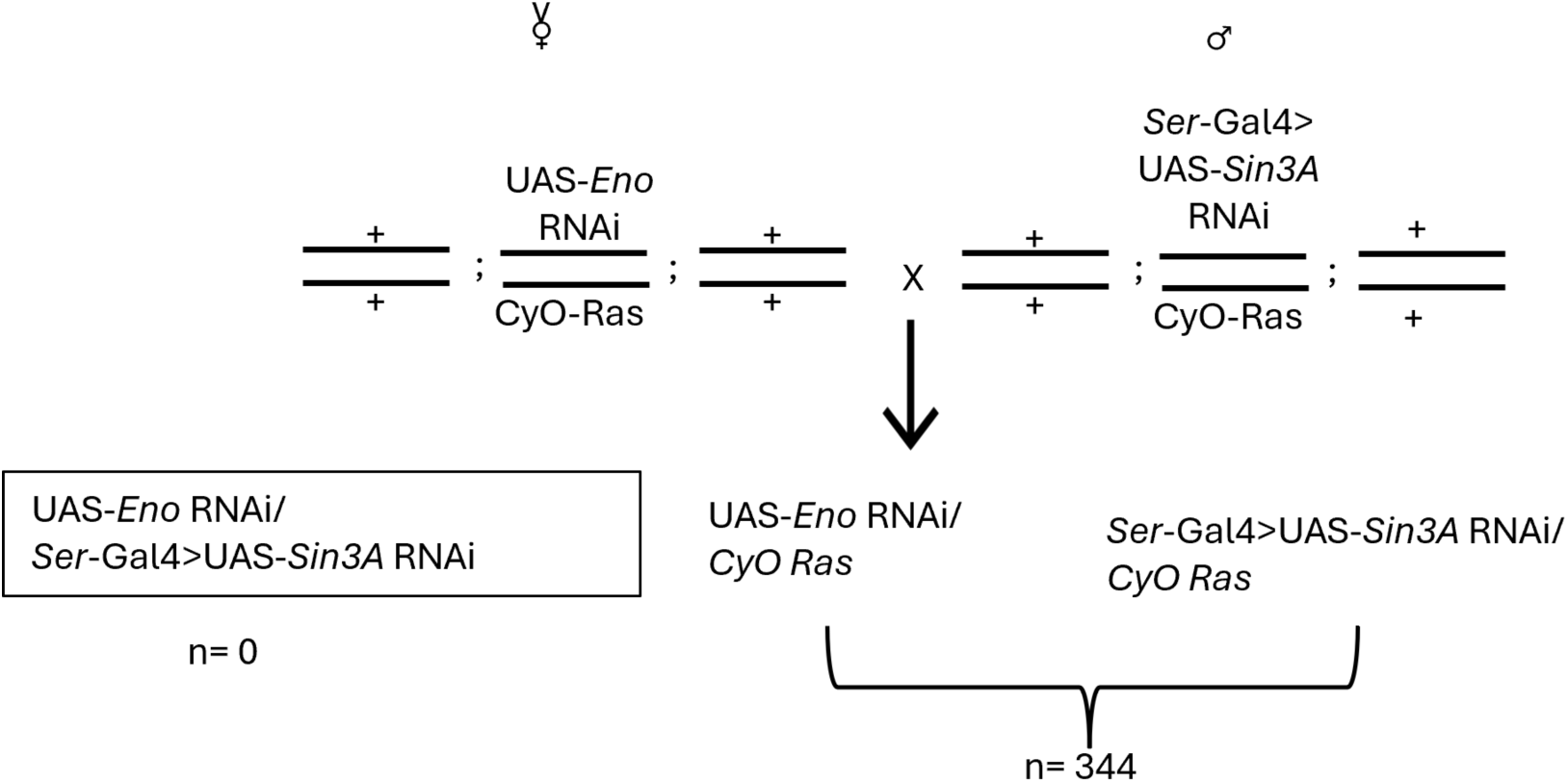
Fly crosses that were done to obtain the *Eno* and *Sin3A* double knockdown.

**Supplementary Fig. 2.**
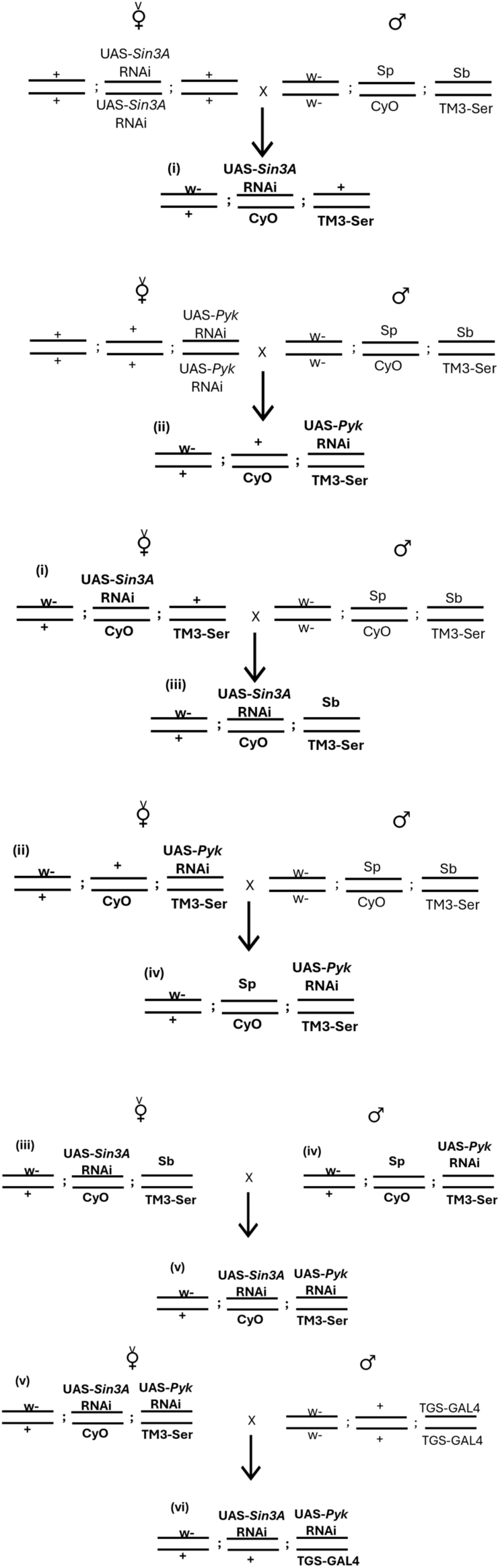
Fly crosses were performed to obtain double-knockdown flies of *Pyk*, *Eno*, *Pfk*, and *Sin3A* that carry TGS-GAL4.

**Supplementary Fig. 3.**
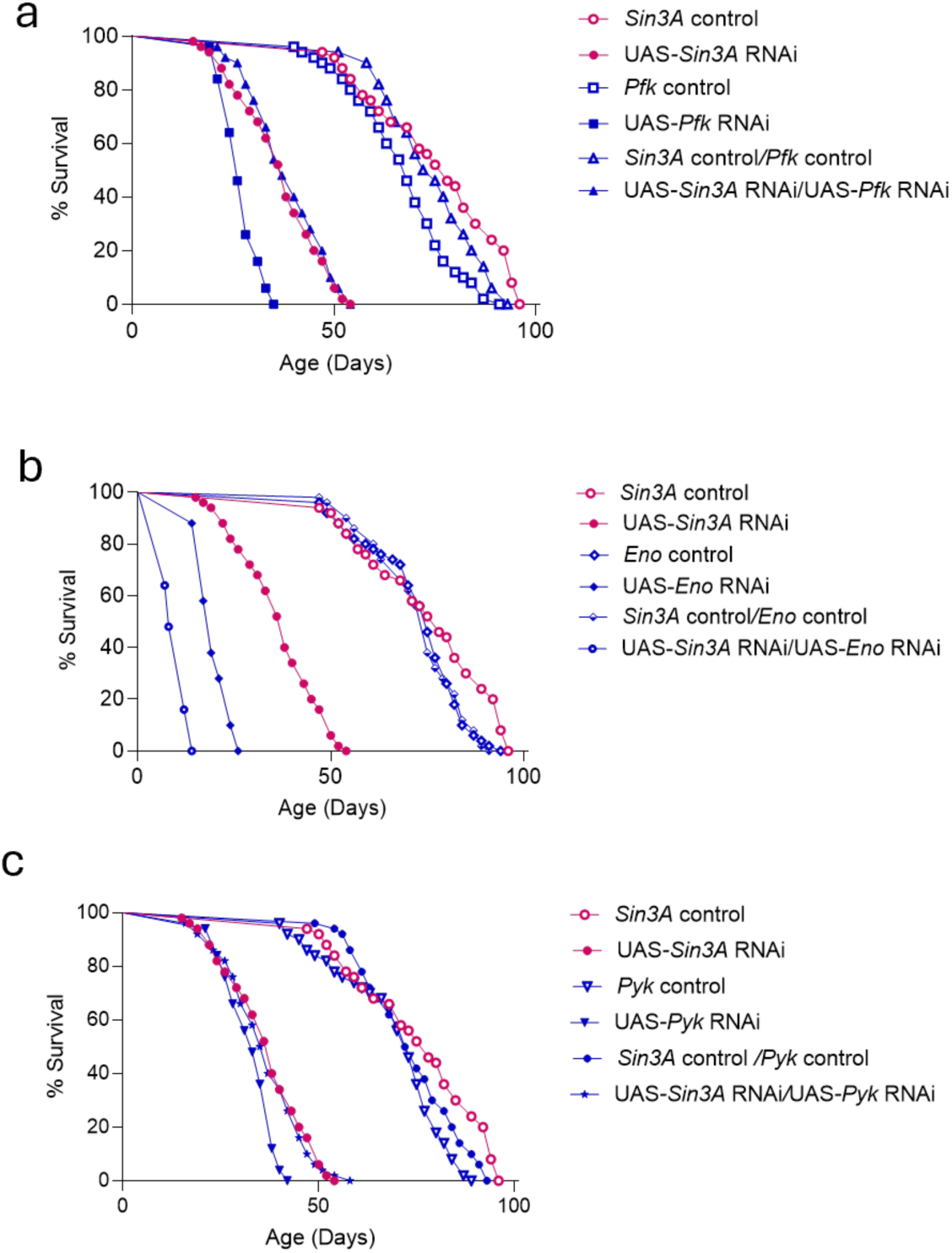
Survival curves of adult male Drosophila under the following conditions: *Sin3A* knockdown alone; *Pyk*, *Eno*, or *Pfk* knockdown alone; and simultaneous knockdown of *Sin3A* with *Pyk*, *Eno*, or *Pfk*.

**Supplementary Fig. 4.**
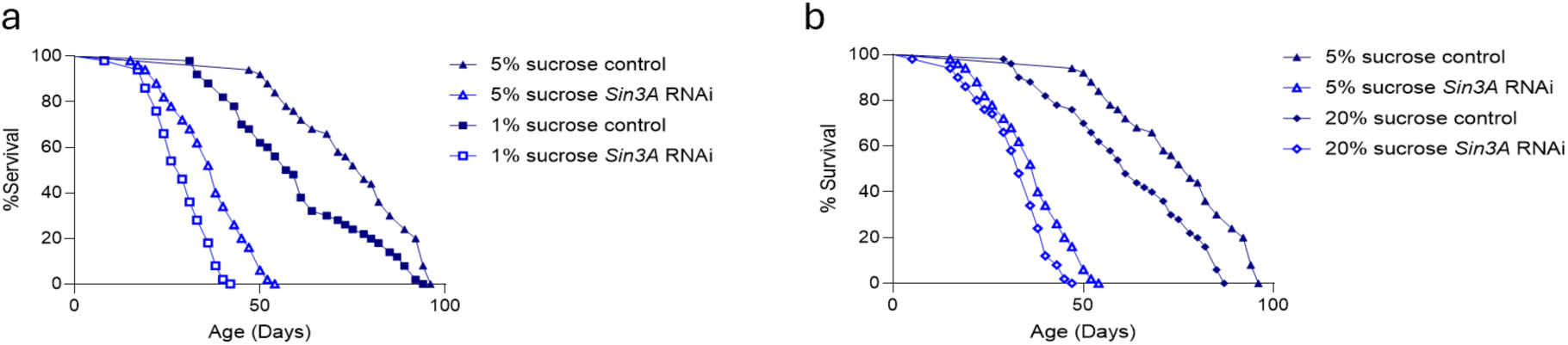
Survival curves of adult male Drosophila reared on food containing 1%, 5%, or 20% (w/v) sucrose.

## References

Adams GE, Chandru A, Cowley SM. 2018. Co-repressor, co-activator and general transcription factor: the many faces of the Sin3 histone deacetylase (HDAC) complex. Biochem J. 475(24):3921–3932. 10.1042/BCJ20170314

Alves-Filho JC, Pålsson-McDermott EM. 2016. Pyruvate Kinase M2: A Potential Target for Regulating Inflammation. Front Immunol. 7 [accessed 2026 Jan 10]. https://www.frontiersin.org/journals/immunology/articles/10.3389/fimmu.2016.00145/full. 10.3389/fimmu.2016.00145

Bannister AJ, Kouzarides T. 2011. Regulation of chromatin by histone modifications. Cell Res. 21(3):381–395. 10.1038/cr.2011.22

Barnes VL et al. 2014. SIN3 is critical for stress resistance and modulates adult lifespan. Aging. 6(8):645–660. 10.18632/aging.100684

Bartolomé A et al. 2022. An Overfeeding-Induced Obesity Mouse Model Reveals Necessity for Sin3a in Postnatal Peak β-Cell Mass Acquisition. Diabetes. 71(11):2395–2401. 10.2337/db22-0306

Bishop JG, Corces VG. 1990. The nucleotide sequence of a *Drosophila melanogaster* enolase gene. Nucleic Acids Res. 18(1):191–191. 10.1093/nar/18.1.191

Carthew RW. 2021. Gene Regulation and Cellular Metabolism: An Essential Partnership. Trends Genet. 37(4):389–400. 10.1016/j.tig.2020.09.018

Chandegra B, Tang JLY, Chi H, Alic N. 2017. Sexually dimorphic effects of dietary sugar on lifespan, feeding and starvation resistance in Drosophila. Aging. 9(12):2521–2528. 10.18632/aging.101335

Chaubal A, Pile LA. 2018. Same agent, different messages: insight into transcriptional regulation by SIN3 isoforms. Epigenetics Chromatin. 11(1):17. 10.1186/s13072-018-0188-y

Chintapalli VR, Wang J, Dow JAT. 2007. Using FlyAtlas to identify better Drosophila melanogaster models of human disease. Nat Genet. 39(6):715–720. 10.1038/ng2049

Cowley SM et al. 2005. The mSin3A chromatin-modifying complex is essential for embryogenesis and T-cell development. Mol Cell Biol. 25(16):6990–7004. 10.1128/MCB.25.16.6990-7004.2005

van Dam E et al. 2020. Sugar-Induced Obesity and Insulin Resistance Are Uncoupled from Shortened Survival in Drosophila. Cell Metab. 31(4):710–725.e7. 10.1016/j.cmet.2020.02.016

Dannenberg J-H et al. 2005. mSin3A corepressor regulates diverse transcriptional networks governing normal and neoplastic growth and survival. Genes Dev. 19(13):1581–1595. 10.1101/gad.1286905

Díaz-Ramos À, Roig-Borrellas A, García-Melero A, López-Alemany R. 2012. α-Enolase, a Multifunctional Protein: Its Role on Pathophysiological Situations. BioMed Res Int. 2012(1):156795. 10.1155/2012/156795

Donati S, Sander T, Link H. 2018. Crosstalk between transcription and metabolism: how much enzyme is enough for a cell? WIREs Syst Biol Med. 10(1):e1396. 10.1002/wsbm.1396

Dung VM et al. 2018. Neuron-specific knockdown of Drosophila PDHB induces reduction of lifespan, deficient locomotive ability, abnormal morphology of motor neuron terminals and photoreceptor axon targeting. Exp Cell Res. 366(2):92–102. 10.1016/j.yexcr.2018.02.035

García M et al. 2009. Phosphofructo-1-Kinase Deficiency Leads to a Severe Cardiac and Hematological Disorder in Addition to Skeletal Muscle Glycogenosis. PLOS Genet. 5(8):e1000615. 10.1371/journal.pgen.1000615

Giovannetti M et al. 2024. SIN-3 transcriptional coregulator maintains mitochondrial homeostasis and polyamine flux. iScience. 27(5) [accessed 2025 Aug 19]. https://www.cell.com/iscience/abstract/S2589-0042(24)01011-3. 10.1016/j.isci.2024.109789

Graveley B. 2010. The developmental transcriptome of Drosophila melanogaster. Genome Biol. 11(1):I11. 10.1186/gb-2010-11-s1-i11

Grüning N-M, Lehrach H, Ralser M. 2010. Regulatory crosstalk of the metabolic network. Trends Biochem Sci. 35(4):220–227. 10.1016/j.tibs.2009.12.001

Grzenda A, Lomberk G, Zhang J-S, Urrutia R. 2009. Sin3. Biochim Biophys Acta. 1789(0):443–450. 10.1016/j.bbagrm.2009.05.007

Heidarian Y et al. 2024. Metabolomic analysis of Drosophila melanogaster larvae lacking pyruvate kinase. G3 GenesGenomesGenetics. 14(1):jkad228. 10.1093/g3journal/jkad228

Hunt LC, Demontis F. 2022. Age-Related Increase in Lactate Dehydrogenase Activity in Skeletal Muscle Reduces Life Span in *Drosophila* Le Couteur DG, editor. J Gerontol Ser A. 77(2):259–267. 10.1093/gerona/glab260

Janke R, Dodson AE, Rine J. 2015. Metabolism and Epigenetics. Annu Rev Cell Dev Biol. 31(Volume 31, 2015):473–496. 10.1146/annurev-cellbio-100814-125544

Jans K et al. 2024. Dietary sucrose determines the regulatory activity of lithium on gene expression and lifespan in Drosophila melanogaster. Aging. 16(11):9309–9333. 10.18632/aging.205933

Jenkins CM, Yang J, Sims HF, Gross RW. 2011. Reversible High Affinity Inhibition of Phosphofructokinase-1 by Acyl-CoA. J Biol Chem. 286(14):11937–11950. 10.1074/jbc.M110.203661

Konwar C et al. 2022. SIN-3 functions through multi-protein interaction to regulate apoptosis, autophagy, and longevity in Caenorhabditis elegans. Sci Rep. 12(1) [accessed 2025 July 8]. https://www.nature.com/articles/s41598-022-13864-0. 10.1038/s41598-022-13864-0

Konwar C, Maini J, Saluja D. 2024. Understanding Longevity: SIN-3 and DAF-16 Revealed as Independent Players in Lifespan Regulation. J Gerontol Ser A. 79(9):glae160. 10.1093/gerona/glae160

Kouzarides T. 2007. Chromatin modifications and their function. Cell. 128(4):693–705. 10.1016/j.cell.2007.02.005

Liu M et al. 2020. A complex interplay between SAM synthetase and the epigenetic regulator SIN3 controls metabolism and transcription. J Biol Chem. 295(2):375–389. 10.1074/jbc.ra119.010032

Liu M, Pile LA. 2017. The Transcriptional Corepressor SIN3 Directly Regulates Genes Involved in Methionine Catabolism and Affects Histone Methylation, Linking Epigenetics and Metabolism. J Biol Chem. 292(5):1970–1976. 10.1074/jbc.m116.749754

Locasale JW. 2018. New concepts in feedback regulation of glucose metabolism. Curr Opin Syst Biol. 8:32–38. 10.1016/j.coisb.2017.11.005

Lushchak OV et al. 2014. Specific Dietary Carbohydrates Differentially Influence the Life Span and Fecundity of Drosophila melanogaster. J Gerontol Ser A. 69(1):3–12. 10.1093/gerona/glt077

Ma Z et al. 2018. Epigenetic drift of H3K27me3 in aging links glycolysis to healthy longevity in Drosophila. eLife. 7 [accessed 2025 July 8]. https://elifesciences.org/articles/35368. 10.7554/elife.35368

Magwere T, Chapman T, Partridge L. 2004. Sex Differences in the Effect of Dietary Restriction on Life Span and Mortality Rates in Female and Male Drosophila Melanogaster. J Gerontol A Biol Sci Med Sci. 59(1):B3–B9. 10.1093/gerona/59.1.b3

Mitra A et al. 2021. Soft repression: Subtle transcriptional regulation with global impact. BioEssays News Rev Mol Cell Dev Biol. 43(2):e2000231. 10.1002/bies.202000231

Mitra A et al. 2022. Isoforms of the transcriptional cofactor SIN3 differentially regulate genes necessary for energy metabolism and cell survival. Biochim Biophys Acta BBA - Mol Cell Res. 1869(10):119322. 10.1016/j.bbamcr.2022.119322

Nasmyth K, Stillman D, Kipling D. 1987. Both positive and negative regulators of HO transcription are required for mother-cell-specific mating-type switching in yeast. Cell. 48(4):579–587. 10.1016/0092-8674(87)90236-4

Nunes RD et al. 2016. Unique PFK regulatory property from some mosquito vectors of disease, and from Drosophila melanogaster. Parasit Vectors. 9(1):107. 10.1186/s13071-016-1391-y

Oka M et al. 2021. Increasing neuronal glucose uptake attenuates brain aging and promotes life span under dietary restriction in Drosophila. iScience. 24(1):101979. 10.1016/j.isci.2020.101979

Osterwalder T, Yoon KS, White BH, Keshishian H. 2001. A conditional tissue-specific transgene expression system using inducible GAL4. Proc Natl Acad Sci U S A. 98(22):12596–12601. 10.1073/pnas.221303298

Pandey R, Sharma M, Saluja D. 2018. SIN-3 as a key determinant of lifespan and its sex dependent differential role on healthspan in Caenorhabditis elegans. Aging. 10(12):3910–3937. 10.18632/aging.101682

Pennetta G, Pauli D. 1998. The Drosophila Sin3 gene encodes a widely distributed transcription factor essential for embryonic viability. Dev Genes Evol. 208(9):531–536. 10.1007/s004270050212

Pessa JC, Joutsen J, Sistonen L. 2024. Transcriptional reprogramming at the intersection of the heat shock response and proteostasis. Mol Cell. 84(1):80–93. 10.1016/j.molcel.2023.11.024

Pile LA, Schlag EM, Wassarman DA. 2002. The SIN3/RPD3 Deacetylase Complex Is Essential for G2 Phase Cell Cycle Progression and Regulation of SMRTER Corepressor Levels. Mol Cell Biol. 22(14):4965–4976. 10.1128/MCB.22.14.4965-4976.2002

Pile LA, Spellman PT, Katzenberger RJ, Wassarman DA. 2003. The SIN3 Deacetylase Complex Represses Genes Encoding Mitochondrial Proteins: IMPLICATIONS FOR THE REGULATION OF ENERGY METABOLISM *. J Biol Chem. 278(39):37840–37848. 10.1074/jbc.M305996200

Poyatos JF. 2020. Genetic buffering and potentiation in metabolism Ma S, editor. PLOS Comput Biol. 16(9):e1008185. 10.1371/journal.pcbi.1008185

Robert VJ et al. 2023. SIN-3 acts in distinct complexes to regulate the germline transcriptional program in Caenorhabditis elegans. Development. 150(21):dev201755. 10.1242/dev.201755

Rodríguez-Vázquez M et al. 2024. Fat body glycolysis defects inhibit mTOR and promote distant muscle disorganization through TNF-α/egr and ImpL2 signaling in Drosophila larvae. EMBO Rep. 25(10):4410–4432. 10.1038/s44319-024-00241-3

Rovenko BM et al. 2015. High sucrose consumption promotes obesity whereas its low consumption induces oxidative stress in Drosophila melanogaster. J Insect Physiol. 79:42–54. 10.1016/j.jinsphys.2015.05.007

Saha N, Liu M, Gajan A, Pile LA. 2016. Genome-wide studies reveal novel and distinct biological pathways regulated by SIN3 isoforms. BMC Genomics. 17(1) [accessed 2025 July 8]. https://bmcgenomics.biomedcentral.com/articles/10.1186/s12864-016-2428-5. 10.1186/s12864-016-2428-5

Sharma M, Pandey R, Saluja D. 2018. ROS is the major player in regulating altered autophagy and lifespan in sin-3 mutants of C. elegans. Autophagy. 14(7):1239–1255. 10.1080/15548627.2018.1474312

Sharma V, Swaminathan A, Bao R, Pile LA. 2008. Drosophila SIN3 is required at multiple stages of development. Dev Dyn Ok Publ Am Assoc Anat. 237(10):3040–3050. 10.1002/dvdy.21706

Soukar I et al. 2025. The histone modification regulator, SIN3, plays a role in the cellular response to changes in glycolytic flux. PLOS ONE. 20(11):e0335411. 10.1371/journal.pone.0335411

Soukar I, Mitra A, Pile LA. 2023. Analysis of the chromatin landscape and RNA polymerase II binding at SIN3-regulated genes. Biol Open. 12(11):bio060026. 10.1242/bio.060026

Sternberg PW, Stern MJ, Clark I, Herskowitz I. 1987. Activation of the yeast HO gene by release from multiple negative controls. Cell. 48(4):567–577. 10.1016/0092-8674(87)90235-2

Strilbytska O et al. 2020. Dietary sucrose defines lifespan and metabolism in Drosophila. Ukr Biochem J. 92(5):97–105. 10.15407/ubj92.05.097

Swaminathan A et al. 2012. Identification of Genetic Suppressors of the Sin3A Knockdown Wing Phenotype. PLOS ONE. 7(11):e49563. 10.1371/journal.pone.0049563

Tanner LB et al. 2018. Four Key Steps Control Glycolytic Flux in Mammalian Cells. Cell Syst. 7(1):49–62.e8. 10.1016/j.cels.2018.06.003

Tennessen JM et al. 2014. Coordinated Metabolic Transitions During Drosophila Embryogenesis and the Onset of Aerobic Glycolysis. G3 GenesGenomesGenetics. 4(5):839–850. 10.1534/g3.114.010652

Volkenhoff A et al. 2015. Glial Glycolysis Is Essential for Neuronal Survival in Drosophila. Cell Metab. 22(3):437–447. 10.1016/j.cmet.2015.07.006

Waller TJ, Collins CA, Dus M. 2025. Pyruvate kinase deficiency links metabolic perturbations to neurodegeneration and axonal protection. Mol Metab. 98:102187. 10.1016/j.molmet.2025.102187

Wang C et al. 2024. Analysis of phosphofructokinase-1 activity as affected by pH and ATP concentration. Sci Rep. 14(1):21192. 10.1038/s41598-024-72028-4

Wong KKL, Liao JZ, Verheyen EM. 2019. A positive feedback loop between Myc and aerobic glycolysis sustains tumor growth in a *Drosophila* tumor model. eLife. 8:e46315. 10.7554/elife.46315

Xin B, Rohs R. 2018. Relationship between histone modifications and transcription factor binding is protein family specific. Genome Res. 28(3):321–333. 10.1101/gr.220079.116

Yang X et al. 2020. Coregulator Sin3a Promotes Postnatal Murine β-Cell Fitness by Regulating Genes in Ca2+ Homeostasis, Cell Survival, Vesicle Biosynthesis, Glucose Metabolism, and Stress Response. Diabetes. 69(6):1219–1231. 10.2337/db19-0721

Yu X, Li S. 2024. Specific regulation of epigenome landscape by metabolic enzymes and metabolites. Biol Rev. 99(3):878–900. 10.1111/brv.13049

Zhang L et al. 2018. Revealing transcription factor and histone modification co-localization and dynamics across cell lines by integrating ChIP-seq and RNA-seq data. BMC Genomics. 19(10):914. 10.1186/s12864-018-5278-5

